# Visible Light Optical Coherence Tomography Reveals Aging at the Retinal Pigment Epithelium-Bruch’s Membrane Interface

**DOI:** 10.64898/2026.03.18.712487

**Authors:** Ruoyu Meng, Rachel C. Kenney, Moning Pan, Alok K. Gupta, Yasha S. Modi, Pooja Chauhan, Christine A. Curcio, Vivek J. Srinivasan

## Abstract

Landmark histological studies have shown that as the retina ages, lipids and other debris accumulate within Bruch’s membrane (BM) and in spaces introduced between BM and the retinal pigment epithelium (RPE). These deposits grow with age, increasing the risk of age-related macular degeneration (AMD), the leading cause of irreversible vision loss for older adults globally. Current *in vivo* imaging lacks specificity to study BM and the important spaces at the RPE/BM interface in living human eyes, while histological techniques suffer from processing artifacts that distort photoreceptors. Here we employ visible light Optical Coherence Tomography (OCT), with 1 micrometer depth resolution, to quantitatively analyze these tissues in living eyes. We identify age-related changes in a human cohort without retinal pathology: thickening and loss of contrast of the hyper-reflective BM band, and thickening of the RPE together with the sub-RPE basal laminar space (RPE+sBL). Both forms of thickening were locally coupled depending on eccentricity, suggesting related biosynthetic mechanisms. A thicker BM and RPE+sBL were locally associated with anomalies in the overlying photoreceptors. Thus, sub-clinical changes in aging eyes detected by visible light OCT resemble early versions of deposits found in AMD. Visible light OCT depicts the relationship between RPE+sBL, BM, and photoreceptors in aging, holding promise to precisely and non-invasively grade ocular phenotypes ranging from normal aging to early AMD.

**Significance Statement:** As humans age, lipids and other debris deposit in Bruch’s membrane (BM) of the human eye (1) and spaces introduced between the retinal pigment epithelium (RPE) and BM. These deposition processes are linked to the eventual development of age-related macular degeneration (AMD), the leading blinding disease amongst older adults. Here we investigate these deposits with visible light OCT imaging in living human subjects without overt retinal pathology. Whereas early RPE/BM deposits were previously assessed only in donor eyes *postmortem* via preparations that distort photoreceptors, our results shed light on these AMD precursors and overlying photoreceptor changes in living human eyes.

## Introduction

Maintaining quality of life in an aging society requires management of age-related diseases, including cardiometabolic disease, cancer, and dementia. Age-related macular degeneration (AMD) is the leading cause of irreversible vision loss for older adults (2). Moreover, AMD is a truly global health concern, with a disease burden that varies by country, depending on race, genetics, income, age, and risk factors including smoking and diet (3). AMD includes an atherosclerosis-like, pro-inflammatory progression (4, 5) where lipids and other debris accumulate with age, first within Bruch’s membrane (BM), and eventually, in the spaces adjacent to the basal lamina of the retinal pigment epithelium (RPE). In AMD, these deposits, observed clinically as drusen when large enough, can promote inflammation as well as lead to retinal hypoxia and consequent macular neovascularization (6). Eventually, AMD can result in atrophy, that is, death of the RPE and choriocapillaris microvasculature and loss of photoreceptors they support. While neovascular AMD can be managed medically, geographic atrophy (GA), the end stage of dry AMD (i.e. AMD without neovascularization), remains challenging to treat effectively. This remains true even with dietary recommendations (7), and more recently, complement inhibitor treatments (8), which modestly slow GA expansion. Based on trial results to date, it is well-accepted that treating at an earlier stage is a promising avenue to improve outcomes (9). Characterizing onset and early progression of AMD is essential in this endeavor.

Optical coherence tomography (OCT) has greatly aided management of AMD by providing cross-sectional imaging of retinal and choroidal architecture with views comparable to low magnification light microscopy of histological sections. OCT has been particularly helpful in depicting the transition to advanced AMD and monitoring progression thereafter. In neovascular AMD, OCT can reveal and track response of exudation to treatment (10). In dry AMD, serial OCT imaging highlights progression of drusen to atrophy (11–13). Several OCT risk factors for progression to GA have been identified (14, 15).

As age is a prerequisite for AMD, changes in the aging eye are seen as setting the stage for the development of AMD (16). In contrast to intermediate and advanced AMD which display salient pathology, aging and AMD onset are characterized by micrometer-scale deposits in spaces at the RPE/BM interface. Relevant deposits, defined based on histology of donor eyes, are 1) lipids, protein, and debris in 3-layer BM (1, 17–19), 2) basal linear deposits (BLinD) together with pre-BLinD, its precursor, between the basal lamina of the RPE and the inner collagenous layer of Bruch’s membrane (20, 21), and 3) basal laminar deposits (BLamD), between the RPE plasma membrane and the RPE basal lamina (22–24). These deposits are observed in older donor eyes without AMD, and to a greater degree, in AMD (20, 22, 25, 26), contributing to a continuum of aging-early AMD phenotypes. Changes in RPE cell morphology (27–29), as well as RPE organelle composition (18, 30, 31) have also been studied, and are important to interpret *in vivo* imaging across the human lifespan.

While highly successful for monitoring intermediate and advanced AMD, the current clinical standard OCT, which uses near-infrared (NIR) light, has faced limitations in studying precursor pathologies at the BM/RPE interface. Commercial NIR OCT systems with 5-7 micrometer resolution cannot reveal let alone quantify BM when it is anatomically apposed with the RPE (Though pathological changes may significantly perturb this relationship to the point of BM visibility (32)). Commercial NIR OCT systems do provide a measure of thickness that nominally includes (from inner to outer) the RPE cell body, its basal lamina, sub-RPE basal laminar space, and BM (RPE+sBL+BM). However, questions remain regarding comparisons to histology and consistency across devices and software packages (33). Prototype NIR OCT of aging and early-intermediate AMD, with a depth resolution of 2.7 micrometers comparable to high-end commercial instruments, highlighted BLamD as an early indicator of AMD (34). This work used BM as a reference surface for quantifying BLamD and stated that BM is rarely distinguishable from the RPE in older subjects at central eccentricities (35). Along these lines, even more granular grading of the RPE+sBL and BM in aged eyes for risk of AMD progression in the early stages *in vivo*, could aid development of interventions targeting early AMD.

Though first demonstrated in 2002 (36), over the past decade, visible light OCT (36–39) has emerged as a useful research tool (40–43). With recent technical advances, visible light OCT now provides unparalleled cross-sectional detail in the outer retina and RPE/BM (44–46). In the context of outer retinal aging, the advantages of visible light for OCT are twofold. First, it provides a depth resolution of 1 micrometer, about 3 times finer than the finest resolution commercial NIR OCT systems. Second, shorter wavelengths improve intrinsic OCT contrast for BM near the foveal center (47), distinguishing BM from the RPE+sBL, defined here as the RPE and sub-RPE basal laminar space but excluding the hyper-reflective BM band. Here, we leverage these advantages, together with a customized scan protocol that serves as a fixation target (see **Materials and Methods**), to characterize the photoreceptors, RPE, and BM in aging (46).

## Results

### Nomenclature

This section discusses nomenclature for the 4 major outer retinal bands in standard clinical OCT (**Figure 1**A). Excepting band 1, which is universally accepted as the external limiting membrane (ELM) (48, 49), there is considerable variability in the OCT nomenclature for outer retinal bands 2 to 4. Here we present our naming conventions for bands 2 to band 4. The goal of this work is not to reconcile terminology, and the conclusions of this work do not depend on a particular attribution of outer retinal photoreceptor bands to subcellular structures.

**Figure 1.**
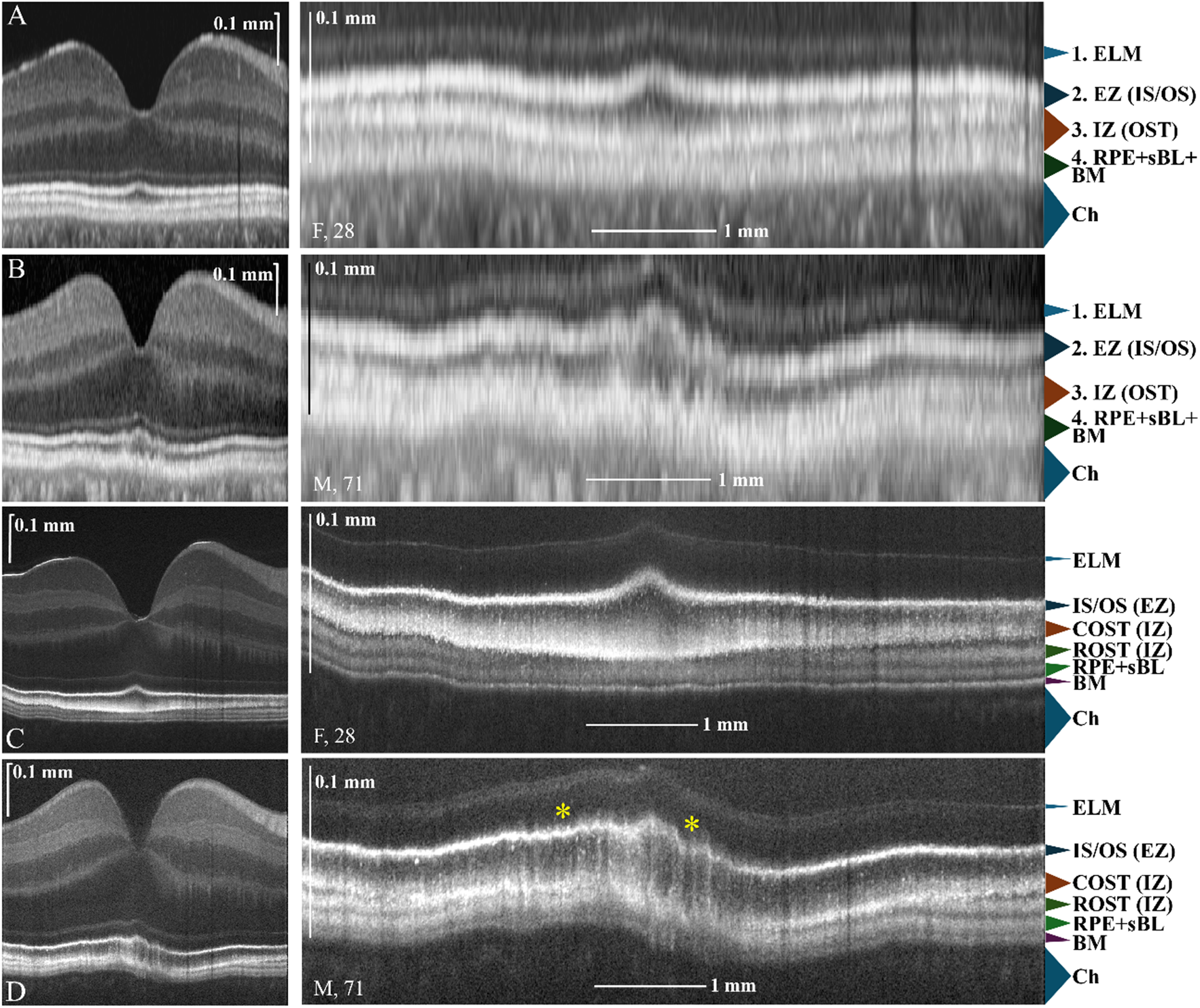
Comparison of Heidelberg Spectralis near-infrared (NIR) OCT (A-B) and prototype visible light OCT (C-D) retinal images in the same eyes: a 28-year-old white female (A,C) and a 71-year-old white male (B,D).In (D), undulations (one of photoreceptor anomalies described in the main text) are observed in the fovea. Layers are named for consistency with literature (see **Nomenclature**). ELM: External Limiting Membrane; EZ: Ellipsoid Zone; IZ: Interdigitation Zone; RPE+sBL: Retinal Pigment Epithelium and sub-RPE basal laminar space; BM: Bruch’s Membrane; Ch: Choroid; IS/OS: Inner Segment/Outer Segment Junction; COST: Cone Outer Segment Tips; ROST: Rod Outer Segment Tips.

#### Band 2

We use the standard clinical terminology, ellipsoid zone (EZ) for Heidelberg Spectralis OCT (48, 49). We use inner segment / outer segment junction (IS / OS) for visible light OCT, providing a parenthetical reference to EZ where appropriate.

#### Band 3

We use the standard clinical terminology, interdigitation zone (IZ), signifying where photoreceptor outer segments and a brush border made of RPE apical processes come together, for Heidelberg Spectralis OCT(48, 49). We use cone outer segment tips (COST) for visible light OCT, providing a parenthetical reference to IZ where appropriate.

#### Band 4

We use the terminology RPE+sBL+BM for Heidelberg Spectralis OCT (48, 49), which is consistent with consensus terminology. In visible light OCT, we observe two bands within band 4 under the central bouquet. The first band within band 4 is the RPE cell body and sub-RPE basal laminar space (together, RPE+sBL) which includes highly scattering apical RPE organelles, the basal infoldings and drusen, if present. The grouping of sBL with the RPE is of little consequence in young eyes, but this nomenclature becomes critical as deposits form with age. The second band within band 4 is BM. We use the three-layer nomenclature (16) for anatomical BM, which is taken to comprise the inner collagenous layer, elastic layer, and outer collagenous layer, excluding sBL. Further from the foveal center, a band gradually emerges between band 3 and the RPE+sBL, which we call rod OST (ROST). This also appears to correspond to RPE2 in a recent publication using 3 micrometer depth resolution NIR OCT (50). In the future, a detailed examination of ROST to ascertain apical process contributions would be helpful (e.g., with validated autofluorescence imaging for melanin (51)). In the absence of such a study, for ROST, we provide a parenthetical reference to IZ where appropriate to acknowledge potential contributions of apical processes.

### Visible light OCT imaging of ocular aging

Matched images from a commercial Heidelberg Spectralis OCT of a young (28-year-old female) and aged (71-year-old male) outer retina show that age-related changes are subtle in clinical OCT (**Figure 1**A-B). In the older retina, the ellipsoid zone (EZ) and interdigitation zone (IZ) appear to undulate, with unclear etiology. In the younger retina, visible light OCT revealed an ordered stack of hyper-reflective bands external to the external limiting membrane (ELM) (**Figure 1**C), with 4 bands in the foveal center, including the inner segment / outer segment (IS / OS) junction, a single cone outer segment tips (COST) band, and a sharp and well-delineated RPE and BM. This stack of 4 bands gradually transitions to 5 bands, around the edge of the fovea (0.75 mm eccentricity), thereupon including both a cone and rod outer segment tips (ROST) band. In the older subject (**Figure 1**D), BM is noticeably thickened but remains distinguishable from the RPE, albeit less so than in the younger subject, across most of the image. A meso-reflective space emerges beneath the hyper-reflective RPE centrally. Above the thickened RPE-BM, a single OST band is observed, and the transition from two OST bands to a single OST band occurs more eccentrically than in the young subject. Undulations of the IS / OS and the single OST band are observed over the thickened RPE-BM. Reflectivity in the myoid just external to the ELM is more prominent in the older subject as previously described (52).

First, the comparison in **Figure 1** underscores numerous age-related visible light OCT findings that are not well-appreciated with commercial NIR OCT. Second, the clear eccentricity-dependence of these findings, as well as prevailing models in the literature (16, 52), motivate a topographic analysis, as described next. Based on these results, we performed imaging of a cohort of eyes from optometry and general ophthalmology at the NYU Langone Eye Center without clinical signs of retinal pathology (see **Materials and Methods**). We analyzed RPE+sBL and BM thicknesses and BM contrast (53) (see **Fig. S1**) within the foveola, fovea, parafovea, and perifovea, with 0.35 mm, 1.5 mm, 2.5 mm, and 5.5 mm diameter ranges around the foveal center respectively (54). The rest of the image was classified as near periphery. These divisions, rather than the conventional Early Treatment Diabetic Retinopathy Study (ETDRS) regions, were chosen for multiple reasons: 1) the foveolar anatomy presented peculiar problems for motion correction that warranted presenting foveolar BM data separately, and 2) the plateau of RPE thickness matched the 1.5 mm foveal definition rather than the 1 mm ETDRS central region (**Fig. S2**M). Note that the fovea was defined as an annulus around the nominal rod-free zone, thus excluding the foveola. We chose this definition, which runs counter to common nomenclature where the fovea includes the foveola, so as to report mutually exclusive sub-regions.

### Aging RPE/BM

An approximate model of aging RPE/BM based on the literature is shown for reference (**Figure 2**A-B). Categories of relevant deposits include 1) lipids, protein, and debris in Bruch’s membrane, 2) BLinD, together with pre-BLinD, its precursor, between the basal lamina of the RPE and the inner collagenous layer of Bruch’s membrane, and 3) BLamD, between the RPE plasma membrane and the RPE basal lamina (**Figure 2**B). The basal infoldings of the RPE are believed to disappear with age (55). As there is no consensus on changes in RPE cell height with age (28, 29), none is depicted. As downward protrusions of BM are not clearly observed in visible light OCT images, intercapillary pillars are not depicted. The apical processes should extend further from the RPE cell body but are not drawn to scale.

**Figure 2.**
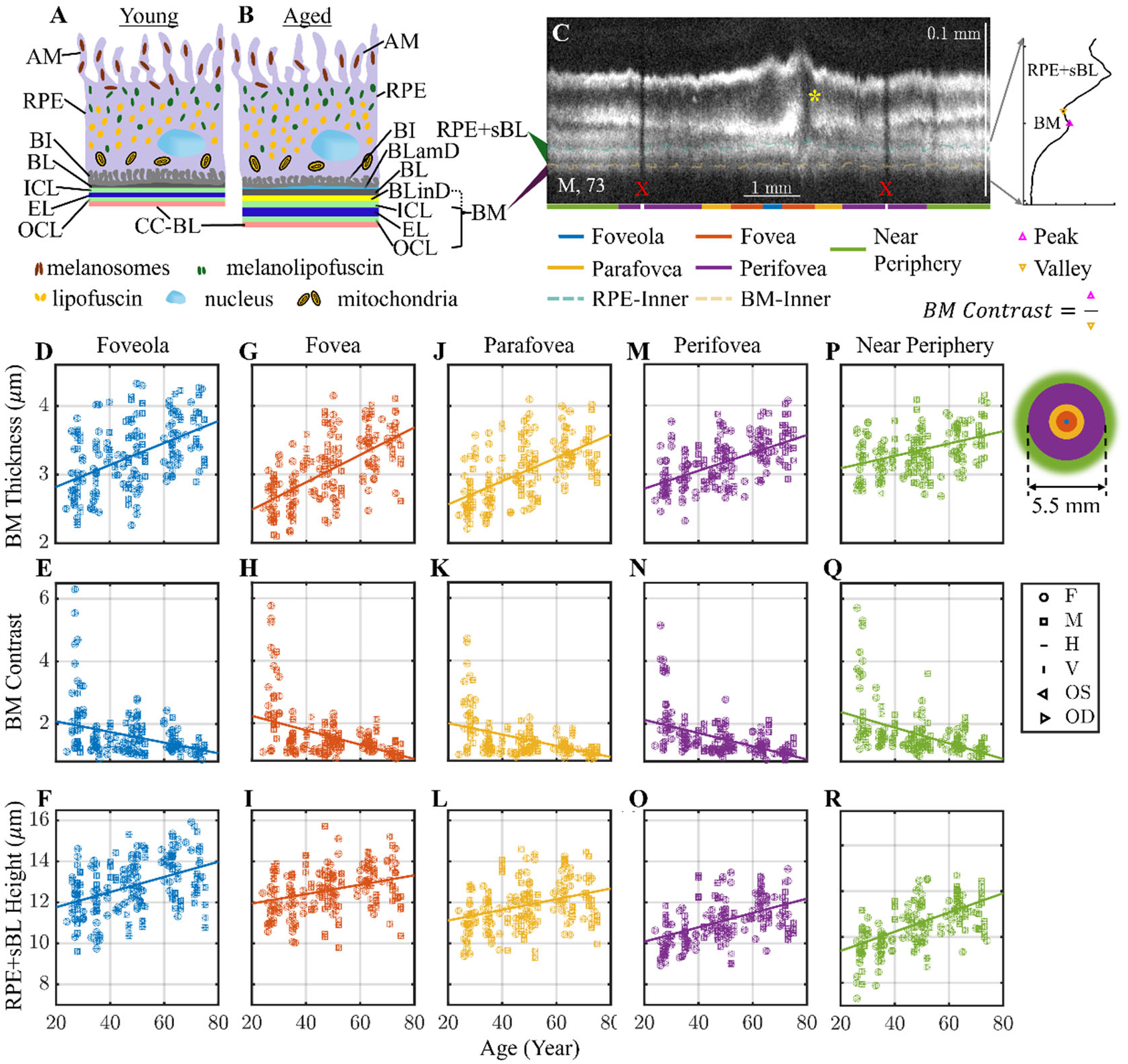
Age-associated changes in RPE+sBL and BM. Pictorial summary of age-related changes from the literature (1, 4, 5, 17–19, 27, 28, 30, 31, 34, 35, 55, 64) in (A) young and (B) aged eyes. With age, there are fewer melanosomes, more lipofuscin and more melanolipofuscin; as well as thicker ICL, EL and OCL due to lipoprotein accumulation. Two additional layers form: basal laminar deposit (BLamD) and basal linear deposit (BLinD). (C) Healthy outer retina of a 73-year-old white male, with 5 sub-regions: foveola, fovea, parafovea, perifovea, and near periphery, color-coded. OCT image is in log scale. The intensity profile of the near periphery subregion is shown on the side panel. BM thickness was defined as the full width at half maximum (FWHM) of the Gaussian fit, and BM contrast was calculated as the peak-over-valley ratio. Note that as per our definition, the fovea excludes the foveola. The segmented boundaries RPE-Inner and BM-Inner define RPE+sBL height. The definitions of BM thickness and contrast are illustrated in the reflectivity profile and further described in **Fig. S1**. The yellow asterisk marks a photoreceptor defect anomaly, showing band 2 elevations and absence of OS/IZ, above a thickened RPE+sBL, which shows different behavior from the shadows caused by blood vessels (red X markers). Across eyes, for each sub-region, BM thickness (D,G,J,M,P) increases with age and modestly decreases with eccentricity; BM contrast (E,H,K,N,Q) decreases with age; RPE+sBL height (F,I,L,O,R) increases with age and decreases with eccentricity. Legend symbols denote female (F), male (M), horizontal (H), vertical (V), left eye (OS), and right eye (OD). IS/OS: Inner Segment/Outer Segment Junction; COST: Cone Outer Segment Tips; ROST: Rod Outer Segment Tips; RPE, retinal pigment epithelium; AM, apical microvilli; BI, basal infoldings; BLamD, basal laminar deposit, BL, basal lamina; BLinD, basal linear deposit, ICL, inner collagenous layer; EL, elastin layer; OCL, outer collagenous layer. Note that the hyperreflective BM is depicted as potentially including early BLinD with a dotted line (see **Discussion**).

### Eccentricity-dependent aging

BM was measured using a previously validated methodology (44) that corrects for RPE multiple scattering (see **Materials and Methods**). BM was found to thicken slightly with eccentricity from the foveola to near-periphery, particularly in younger subjects (**Figure 2**D, G, J, M, P, **Table 1**, **Fig. S3**). Critically, with age, BM thickened (**Figure 2**D, G, J, M, P, **Table 1**) and its contrast (**Figure 2**C) relative to the basal RPE+sBL (**Figure 2**C) declined (**Figure 2**E, H, K, N, Q, **Table 1**). RPE+sBL thinned with eccentricity but thickened with age (**Figure 2**F, I, L, O, R, **Table 1**). RPE+sBL and BM parameters in the foveal sub-region correlate with those of other eccentricities (**Fig. S3**).

**Table 1.** P-values for slopes versus age, intercepts, and slopes versus age were estimated using GEE models for BM thickness, BM contrast, and RPE+sBL height. Significant age-related changes were detected across sub-regions.

| *Changes* | *Model parameters* | *Foveola* | *Fovea* | *Parafovea* | *Perifovea* | *Near Periphery* |
| --- | --- | --- | --- | --- | --- | --- |
| *BM Thickness (μm)* | P-value | $<1 \times 10^{-6}$ | $<1 \times 10^{-6}$ | $<1 \times 10^{-6}$ | $<1 \times 10^{-6}$ | $5 \times 10^{-6}$ |
|  | Intercept (μm) | 2.50 | 2.09 | 2.22 | 2.53 | 2.91 |
|  | Slope (μm/yr) | 0.016 | 0.020 | 0.017 | 0.013 | 0.009 |
| *BM Contrast* | P-Value | $6.0 \times 10^{-3}$ | $6.6 \times 10^{-4}$ | $2.6 \times 10^{-4}$ | $9.5 \times 10^{-5}$ | $2.6 \times 10^{-4}$ |
|  | Intercept | 2.42 | 2.69 | 2.36 | 2.55 | 2.88 |
|  | Slope (1/yr) | -0.017 | -0.023 | -0.018 | -0.021 | -0.025 |
| *RPE+sBL Height (μm)* | P-Value | $1.9 \times 10^{-5}$ | $1.3 \times 10^{-3}$ | $6.6 \times 10^{-4}$ | $<1 \times 10^{-6}$ | $<1 \times 10^{-6}$ |
|  | Intercept (μm) | 11.02 | 10.19 | 11.48 | 10.60 | 8.55 |
|  | Slope (μm/yr) | 0.037 | 0.021 | 0.023 | 0.026 | 0.042 |

### Eccentricity-dependent relationship between RPE+sBL and BM

A tight coupling between RPE+sBL height and BM thickness, defined as a significant slope in a linear fit, was observed (**Figure 3**). As might be expected from the topographic dependence of the RPE thickness, in particular, this correlation coefficient varied with eccentricity from -0.02 to 0.40 (**Table 2**). When data were pooled across eccentricities, the correlation coefficient of RPE+sBL and BM dropped to 0.09, far less than some individual eccentricities.

**Figure 3.**
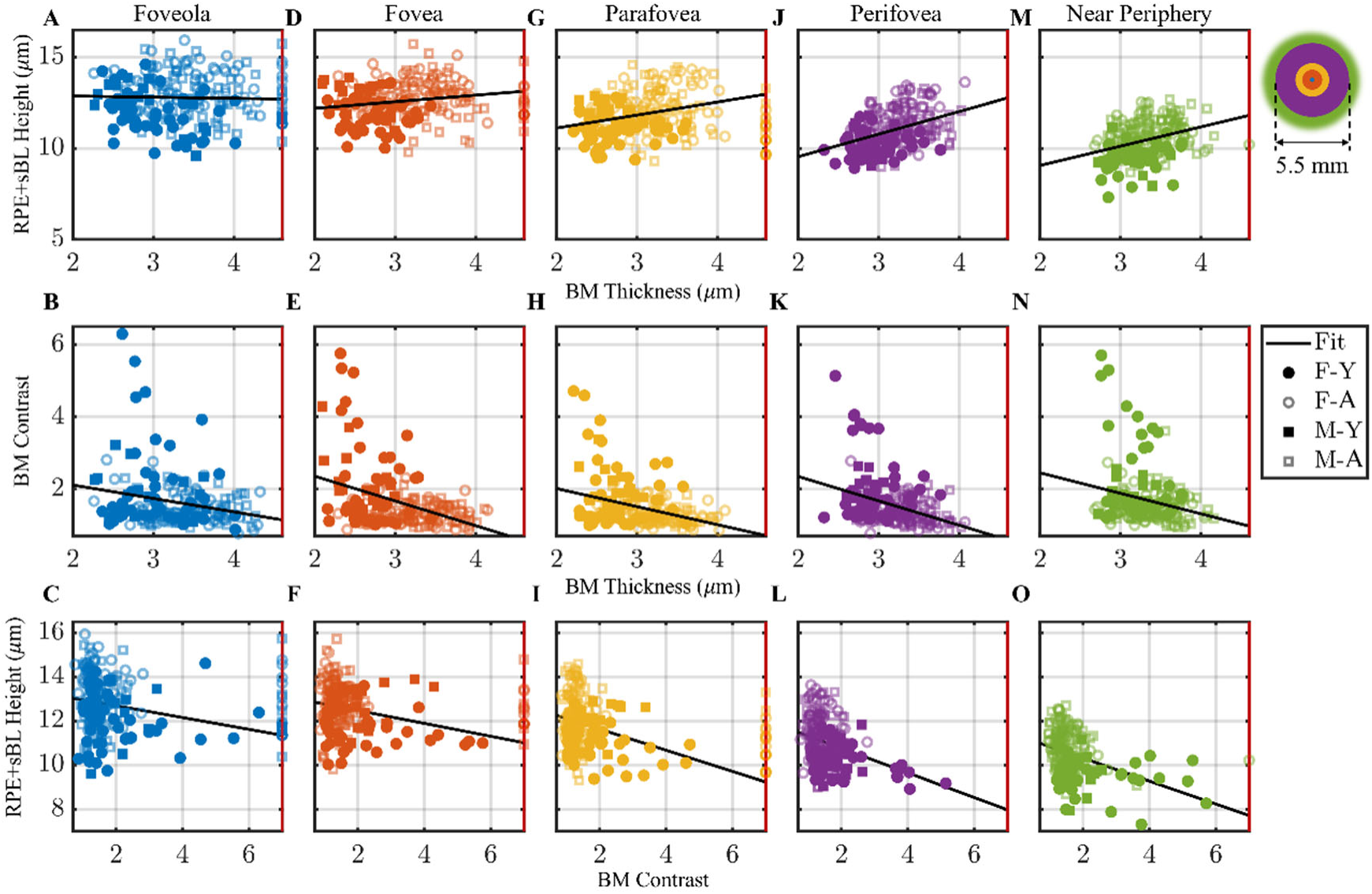
Eccentricity-dependent coupling of (A, D, G, J, M) RPE+sBL height and BM thickness;(B, E, H, K, N) BM contrast and BM thickness; (C, F, I, L, O) RPE+sBL height and BM contrast for sub-regions. All regions had a statistically significant correlation (p<0.05). Subjects under 40 years were classified as younger (Y, <40 years; hollow marker), and those 40 years or older were classified as aged (A, ≥40 years; solid marker). As BM was not detectable in 4.1% of the sub-regions, RPE+sBL heights for these cases are reported as scatterers on the right y-axis.

**Table 2.**
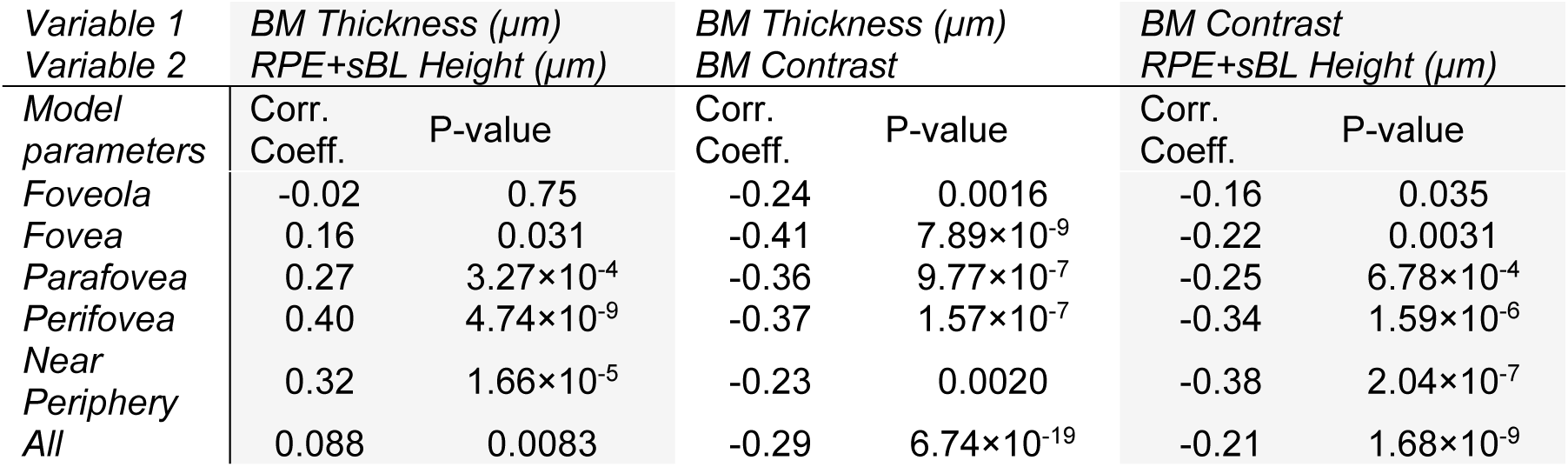
Pairwise Pearson correlation coefficients and p-values between BM thickness, BM contrast, and RPE+sBL height. Note that because of anatomical variations, correlations stratified by sub-region are generally stronger than correlations pooling across sub-regions (last row).

### Photoreceptor anomalies are associated with thicker and less distinguishable BM

Finally, sub-regions of all images were graded for photoreceptor anomalies (defined as focal elevations, undulations, or abnormally absent photoreceptor bands that cannot be explained by directional effects) by consensus of two expert graders (R.M. and V.J.S.). Cases of vessel shadowing were not counted as anomalies (**Figure 2**C, **Figure 4**A, red Xs). BM thickness, RPE+sBL thickness, and BM contrast were compared in sub-regions with and without photoreceptor anomalies (**Figure 4**A-B). We found that BM was significantly thicker in the fovea and parafovea underneath photoreceptor anomalies (**Figure 4**C, **Fig. S2**.D and G), versus no anomalies, using GEE models (Table S1). A similar finding followed for RPE+sBL in the perifovea. While in all other sub-regions, BM and RPE+sBL trended thicker and BM contrast trended lower under photoreceptor anomalies (**Figure 4**D-E, Table S1, **Fig. S4**), false discovery rate (FDR)-corrected p-values were not statistically significant.

**Figure 4.**
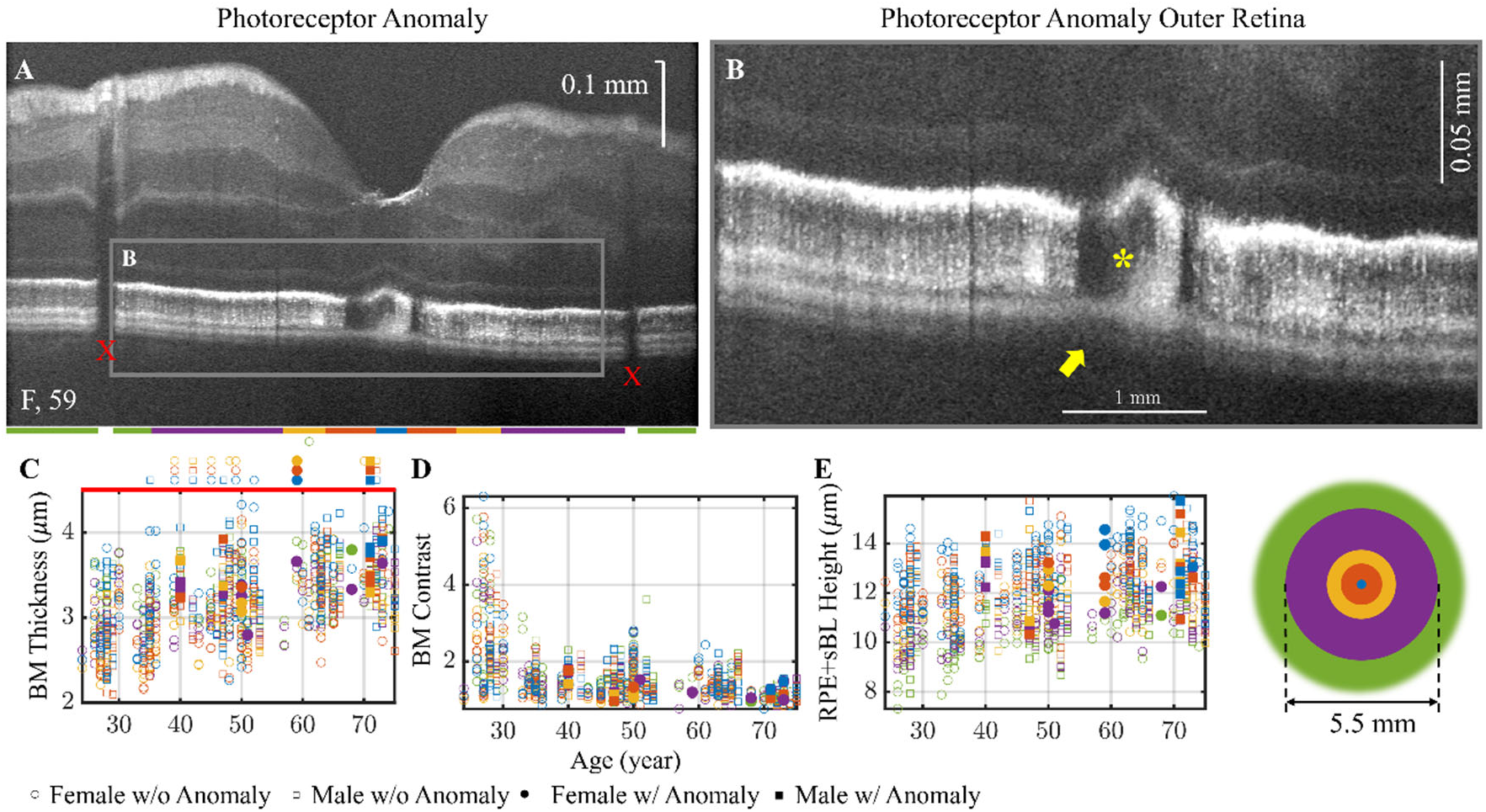
Altered anatomy in sub-regions with photoreceptor anomalies, defined as either abnormal photoreceptor elevations or undulations, or missing photoreceptor bands (excluding vessel shadows marked as red Xs) in a 59-year-old white female. The yellow asterisk marks the photoreceptor defect anomaly, showing absence of OS above a thickened RPE+sBL and weakened BM reflectivity (yellow arrow) leading to lower contrast (A-B). Red X markers mark the shallows resulting from retinal blood vessels. Below photoreceptor anomalies, (C) BM is thicker (**Fig. S2**.A, D, G, J and M); (D) BM contrast is lower (**Fig. S2**.B, E, H, K and N); and (E) RPE+sBL is taller (**Fig. S2**.C, F, I, L and O). BM thickness was found to be a statistically significant predictor of anomalies in the fovea and parafovea, and RPE+sBL in perifovea (**Table S1**). Undetectable BM (shown above the plot in C) was also associated with the presence of photoreceptor anomalies. Sub-regional comparisons from the GEE model are shown in Table S1. Solid and hollow makers denote sub-regions with and without photoreceptor anomalies, respectively.

Sub-regions where BM was undetectable did not provide a BM thickness or contrast. The association of these sub-regions with photoreceptor anomalies, shown above **Figure 4**D, was analyzed separately. Among sub-regions with an anomaly, 9 out of 38 (23.68%) had an undetectable BM, while only 30 out of 906 (3.31%) normal photoreceptor sub-regions had an undetectable BM. A Chi-Squared test yields a ꭕ^2^ of 18.24 and a p-value of 6.32×10^−10^, which indicates that BM is more often undetectable below regions with photoreceptor anomalies.

### Confounding factors

Typically, it is considered challenging to image through the RPE, whose pigmented organelles (melanosomes and melanolipofuscin) are very highly scattering. Though visible light provides some advantages in this regard (47), a critical examination of the validity of the results from the standpoint of optical imaging physics is warranted. In doing so, we consider several potential non-biological confounds that may influence the observed results. The first potential confound is the overall worse image quality observed in older subjects, largely influenced by challenges in maintaining fixation, and age-related media opacities. However, we found that visible light OCT band 2 (IS / OS or EZ inner half width, defined as the distance from the inner half-maximum to the peak, does not change with age (**Fig. S5**). This provides evidence suggesting that worse image quality does not necessarily impair thickness measurements.

A second potential confound is that the local backscattered spectrum could change at the level of BM, for instance, due to absorption and scattering of light in the RPE. If present, this effect would change the local depth resolution of OCT, which depends on the bandwidth of the local interference spectrum. However, we found that the transform-limited resolution of the intensity point spread function, as determined from the spectrum of the basal RPE+sBL and BM, did not vary appreciably with age, and even slightly improved (**Fig. S6**C), remaining well below the nominal 2-4 micrometer thickness of BM (**Fig. S6**). Therefore, changes in the spectrum with age cannot account for thicker BM measurements in older subjects, showing that BM thickening measured by visible light OCT is not a result of worse depth resolution.

A third potential confound arises from the well-known effect that light can scatter multiple times before being detected, leading to appearance of artifacts below highly forward scattering tissues such as blood within vessels (56). Since vessel shadows were excluded from analysis, for this study, the remaining concern is that high RPE multiple scattering could somehow cause a broadening of the BM band in older subjects. The apical peak to basal valley RPE intensity ratio (RPE attenuation), which provides a coarse measure of attenuation due to absorption and scattering, does increase slightly with age (**Fig. S2**). Yet, our method of deriving BM thickness (44) is designed to reduce multiply scattered light and tested in a Monte Carlo simulation (44, 47). Also, the RPE attenuation is greatest in the fovea (**Fig. S2**K) where measured BM is thinnest (**Figure 2**G), a counterexample to this potential confound. Nonetheless, such a counterexample does not disprove that multiple scattering from a thick RPE may impact BM measurements, despite our correction procedure. As a secondary check, the longer wavelength portion of the spectrum increases the contribution of multiple scattering relative to BM backscattering, and yet, scatter-corrected BM thickness does not differ from the shorter wavelength portion of the spectrum where multiple scattering contributions are reduced relative to BM backscattering (**Fig. S7**). This suggests that our method of deriving BM thickness is not biased by variations in multiple scattering.

## Discussion

Aging of the human eye is characterized by a multitude of sub-RPE deposits that have been distinguished histologically, but not yet via *in vivo* imaging. Building on prior efforts (34, 35, 52), here we distinguish and independently quantify BM and RPE+sBL thickening in living human eyes. We first discuss morphometric changes, followed by their association with photoreceptor anomalies, and finally, study limitations and future directions.

### BM thickness

Prior histochemistry and electron microscopy of aged donor eyes has shown that BM undergoes elastic fiber calcification and collagen fiber cross-linking (57), which accompany lipoprotein deposition (1) and leads to overall thickening (19). The observed visible light OCT thickening of our RPE-scatter-corrected BM profile is likely related to these processes. Our BM thickness measurement is designed to be relatively independent of RPE scattering, as validated above.

With age, lipoproteins appear in (3-layer) BM, then between BM and the RPE basal lamina as pre-BLinD, eventually forming BLinD in the same space. To what degree do these deposits contribute to our measured BM thickness? It is reasonable to speculate that, based on proximity, pre-BLinD and early BLinD may be captured in the BM band measurement, as lipoproteins accumulate adjacent to the ICL of 3-layer BM. However, the contribution of advanced BLinD to BM thickness is less likely, because eventually BLinD forms soft drusen (20), which appears as a meso-reflective space distinct from BM. Therefore, we speculate that advanced BLinD and eventual soft drusen would be likely included in our RPE+sBL, not BM, measurement. The potential contribution of BLinD to our BM measurement, if any, requires further study.

Prior studies have noted a doubling of BM thickness with age from age 20-90 years. Across eccentricities, we find the BM band thickens less than this, even when extrapolating from the fit of our cohort to the 20–90-year age range. One potential reason may be the resolution of visible light OCT, which is ideally 0.7 micrometers (intensity FWHM), though the practical resolution is degraded by uncorrected motion, dispersion compensation error, spectral loss (**Fig. S6**), and imperfect registration when averaging. All of these effects place a ‘floor’ on the minimum measurable BM thickness. In addition, we find that reducing the spectral width by about 25% resulted in a thickening of BM of 14.8% at shorter wavelengths and 12.2% at longer wavelengths, showing that resolution is an important determinant of our estimate of BM thickness (7). Another potential reason is that we defined our cohort based on a recent clinical exam to exclude any findings that may indicate AMD. This practice may have removed any borderline AMD cases that would undoubtedly have been included in tissue-level studies that analyze donor eyes. Indeed, the data in prior studies (17, 18) and our data (16) show outliers with a thicker BM amongst aged eyes, leading to a skewed distribution, with some BM thickness values exceeding 5 micrometers. By comparison, we observed no such skew in our *in vivo* measurements, and BM was less than 5 micrometers thick. However, our cohort did not include any subjects above 80 years old due to our inclusion criteria. It is also possible that prior BM histomorphometry included intercapillary pillars (58), or even BLinD or even BLamD, in their measurements of BM. Of note, we did not clearly observe “Roman arch bridge” features suggestive of downward extensions of the outer collagenous zone into intercapillary pillars (59), although they should have been resolvable by visible light OCT, if present. We also had to exclude BM measurements in 4.1% of sub-regions where there was no clear BM profile; this may have effectively excluded regions with a thicker BM if a thicker BM was not detectable. We plan to investigate early AMD cases with visible drusen and pigmentary changes in a future study.

### BM contrast

BM contrast is defined as the ratio of two quantities and is consequently more challenging to interpret than BM thickness. BM contrast reduction implies that with age, the reflectivity of BM is reduced relative to the reflectivity, inclusive of multiple scatter, of the overlying basal RPE and basal laminar space (RPE+sBL) (44). It is conceivable that as BM thickens, it becomes less reflective. However, the denominator of BM contrast, as defined, is the reflectivity depicted in the region immediately inner to BM, which is nominally the basal RPE and infoldings, but also includes multiply scattered light from the apical and middle RPE (47). Therefore, RPE pigmentation and organelle distribution (i.e. the gap between the scattering organelles and BM), as well as the appearance and growth of basal laminar and linear deposits, may also play an important role in BM contrast. Though contrast results are reported, there are many contributing biological factors making interpretation unclear. At a minimum, BM contrast is a useful and concise metric of its visibility in OCT images. We advocate its use in imaging studies of BM moving forward.

### RPE+sBL height

Another feature of this study was the segregation of RPE+sBL, the RPE inclusive of the sub-RPE basal laminar space. The apical border of RPE+sBL is the rise of scattering from intracellular organelles, and the basal border is the inner edge of BM. Our RPE+sBL thickness measurements are generally in line with histology (28, 29, 60). However, our RPE inner boundary is defined by distributions of organelles and therefore may not precisely demarcate the actual RPE cell boundary (61). It is conceivable that a redistribution of organelles within the cell can change our measured RPE cell height, even if the actual apical boundary does not change. Interestingly, commercial NIR OCT systems also provide a nominal RPE+sBL+BM measurement. While some values (62, 63) greatly exceed our RPE cell height (28, 33, 64), suggesting contributions of additional bands apart from RPE+sBL and BM, other studies (65–68) are more in line with ours. Our approach is complementary to prior NIR OCT works that defined a hyporeflective basal RPE+sBL band above BM. The segmentation of a hyporeflective basal band (34, 35) nonetheless remains sensitive to distributions of organelles in the middle RPE cell body as well as RPE multiple scattering, whereas RPE+sBL height depends on the distance from the apical organelles to inner BM. Each approach has its merits.

The eccentricity and age-dependence of RPE+sBL merits detailed discussion. The eccentricity dependence of RPE+sBL is likely driven by eccentricity dependence of RPE cell height (28, 29, 60), particularly in young subjects. We find that RPE+sBL thickens with age across all subregions. By comparison, while age-related RPE+sBL+BM thickening has been reported on commercial NIR OCT systems (66, 68, 69), there is a trend towards central thinning(65, 66). RPE+sBL thickening with age may arise from either changes in the RPE cells themselves, or the sub-RPE basal laminar space, or both. In some subjects, we observed an increase in the hyporeflective region above BM, outer to the hyper-reflective apical RPE (**Figure 1**D). Judging from similarity in appearance to purported basal laminar deposit (BLamD) found in early AMD eyes (34, 35), these are likely BLamD. Notably, in this work, such BLamD is detected in aged eyes of a cohort strictly defined to exclude AMD, a finding supported by numerous histological reports (21, 22). RPE cell height is a large contributor to RPE+sBL and should be accounted for in interpretation. Increases in RPE cell height with age have also been reported (28), though the opposite is reported in other studies (29, 33). Reasons for these discrepancies lie in limitations of studies using human donor eyes. In the histology literature, many measurements were made from RPE of detached retina, without accounting for the apical processes, or on specimens without photo-documentation, obviating quality-control in hindsight. As discussed above, the contribution of BLinD to our BM measurement versus our RPE+sBL measurement also requires further study. Ultimately, distinguishing BLinD and BLamD would be clinically significant given the distinct AMD pathways that these deposits portend (25, 32, 70).

Further insights can be gleaned by comparing RPE+sBL height and BM thickness, which may probe different age-related deposits. When eccentricity is not accounted for, we observe a weak correlation of 0.09 (though still statistically significant) between BM and RPE+sBL. On the other hand, when eccentricity is accounted for, we observe that local correlations between BM and RPE+sBL markedly increase, rising as high as 0.40 in the perifovea. This supports a topographically varying relationship between RPE+sBL and BM. These results are consistent with a sequence where lipids deposit first within BM and eventually in the sub-RPE space, if deposits account for age-related thickening of RPE+sBL. We cannot decipher temporal sequence without longitudinal data, however. Results could also be consistent with a potential sequence where BM thickening leads to thickening in the RPE cell. Note that the correlation between BM and RPE+sBL is weaker in the fovea and foveola. This does not necessarily imply that there is no relationship: indeed, the weaker correlation may be due, in part, to the intrinsically greater variability in foveal RPE cell height in younger subjects (double the standard deviation of peripheral RPE as per **Figure 2**F,R) or challenges with measuring BM accurately in the foveola.

For more granular understanding of age-related changes in RPE+sBL, we analyzed its internal reflectivity (**Fig. S2**). A continuous, rather than segmentation-based, analysis (34), was undertaken because clear discrete, hyporeflective sub-RPE regions were challenging to define in all subjects in our aged cohort. As previously described (44), in young eyes, apical RPE organelles were found to be hyper-reflective, whereas the basal RPE+sBL (which mainly comprises basal RPE since deposits are minimal), extending to BM, was comparatively less reflective. The contributions of lipofuscin and mitochondria in visible light OCT may indeed be reduced by powerful visible light shadowing by apical pigmented organelles in healthy RPE. The most salient age-related change was the decrease in the contrast of BM, as noted already. The next observation was that reflectivity spread into a plateau in the apical RPE in older eyes, suggesting a broader apical-mid RPE distribution of scattering organelles. This is consistent with melanolipofuscin, which, during normal aging (27, 30, 31), replaces melanosomes, which are apically concentrated in young eyes. Critically, melanolipofuscin is more broadly distributed in the apical-mid RPE (27) than melanosomes in the apical RPE, consistent with the observed reflectivity plateau (**Fig. S2**A-E). Importantly, the extent of the hyporeflective portion of the RPE+sBL remained comparable in younger and older eyes on the % RPE+sBL length scale. Thus, across the entire cohort, we can make no conclusive ascription of RPE+sBL thickening with age to either the hyper-reflective (organelles) or the hyporeflective space which extends from basal cell body to BM. However, the fact that we observed deposits strongly resembling BLamD in some subjects (e.g. **Figure 1**), still stands.

### Photoreceptor anomalies

Visible light OCT imaged architecture of photoreceptors, the cells responsible for initiating vision, over locations where BM and RPE+sBL were quantified. This capability surpasses conventional histological methods that measure BM and RPE+sBL, but distort the delicate photoreceptor anatomy. Our *in vivo* imaging showed that under photoreceptor anomalies, BM was more often indistinguishable. Where BM could be quantified, it was significantly thicker in the fovea and parafovea in the presence of photoreceptor anomalies. Similarly, RPE+sBL was significantly thicker in the perifovea in the presence of photoreceptor anomalies. The near periphery was consistent with these findings, but since only one photoreceptor anomaly was found, significance was not assessed (**Fig. S5**). These findings are consistent with deposits contributing to inflammation and impaired clearance of waste which lead to photoreceptor damage. Though anomalies were readily detectable on visible light OCT and often, by Spectralis under the same viewing conditions (e.g. **Figure 1**), photoreceptor anomalies which involved absent IS/OS (EZ) were less than 1 visual degree in lateral extent. It is unknown how these anomalies and associated deposits progress with time, and the visual consequences remain unclear. These questions will be the topic of future studies.

### Summary and future directions

Compared to histochemistry and ultrastructure studies which serve as the gold standard for this work, the non-invasive and non-destructive imaging approach presented here has many advantages. These include the ability to study photoreceptors and underlying deposits without post-mortem distortions introduced by tissue processing, the ability to clinically characterize the cohort prior to imaging, and the potential for longitudinal monitoring. On the other hand, there are many limitations of this work. The first is that it is not possible to study the choriocapillaris, an essential partner of RPE/BM in supporting photoreceptor metabolism, due to limited penetration depth of visible light. Secondly, while BM contrast is improved at shorter wavelengths (47), BM was indistinguishable from the RPE multiple scattering tails in 4.1% of sub-regions, precluding an assessment of thickness. Third, since the basal lamina is approximately 0.15 micrometers thick, images cannot clearly distinguish the sub-RPE basal laminar space from the basal RPE cell body, leading to ambiguities in interpretation (34). Fourth, light backscattering provides little biochemical specificity. Along these lines, new spectroscopic imaging and display methods (71) may eventually help. Fifth, to our knowledge the major ophthalmic manufacturers have not yet initiated visible light OCT commercialization efforts, making it a research tool at the moment. However, visible light OCT findings may complement and guide research efforts with current commercial grade instruments (34, 50).

In summary, this work details aging of the BM, RPE+sBL, and the photoreceptors, moving *in vivo* imaging closer to the level of detail previously achieved in donor eyes. A number of questions arise regarding the nature of these age-related changes, which are now accessible in living subjects. First, how does thickening of BM and RPE+sBL relate to rod-mediated dark adaptation, which assesses transport of vitamin A derivatives from the choriocapillaris to the retina across thickened tissues, and is slowed with age (72, 73). Second, does the considerable variability in Bruch’s and RPE morphology, even at a given age, help to stratify for AMD risk? Third, does BM thickening predict subsequent photoreceptor defects or do they proceed hand-in-hand? Fourth, how do choriocapillaris changes relate to BM and RPE+sBL thickening? These questions can now be addressed with visible light OCT *in vivo*, in conjunction with other conventional modalities, as appropriate. Ultimately, answers to these questions may help to predict AMD progression early, speeding progress toward more effective treatments.

## Materials and Methods

### Visible light OCT system and protocol

A prototype spectral domain visible light OCT system was developed, optimized, and installed at the NYU Langone Eye Center for this study (74). The system uses a grating light valve (GLV) spatial light modulator to shape the incident light (75) to achieve a red light spectrum for alignment and a Gaussian white light spectrum for image acquisition with 1 micrometer axial or depth resolution (0.71 micrometer intensity resolution after incoherent averaging to reduce speckle). The visible light OCT spectrum ranged from 491-642 nm. Compared to an exemplary ultrahigh resolution NIR OCT system with a spectral range of 767-915 nm (76), which has similar OCT hardware specifications to the HighRes Spectralis from Heidelberg Engineering (58, 77), we highlight two important differences. First, though the full-width-at-half-maximum (FWHM) wavelength bandwidth (Δ*λ*) of the visible light OCT system is only ∼25% broader, the shorter visible center wavelength (*λ*_o_) and slightly larger (∼1%) group refractive index (*n*) enables 2.7x finer depth resolution (Δ*z*) as per the OCT resolution equation for a Gaussian spectrum (78):

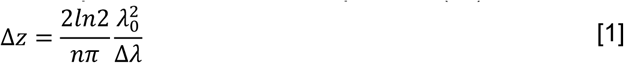

Second, visible light is significantly more impacted by retinal chromophores than NIR light. On the positive side, pigmented organelle absorption and scattering differences at visible wavelengths improve the contrast of BM, particularly near the foveal center (47). On the negative side, due to enhanced visible light absorption by hemoglobin, retinal vessel shadows are stronger, and OCT signal penetration to the choroid is reduced.

The technical challenges with visible light OCT leading to its slow adoption have been extensively discussed in the literature, and many have been addressed (79). One of the challenges that have not been addressed, to our knowledge, is the inability of some patients to fixate on a target during a bright visible raster scan. Here, to address this problem, we design the visible light OCT scan pattern itself to serve as the fixation target. An interleaved horizontal and vertical (X-Y) raster scan pattern served as a fixation target, with ordering of scans chosen to maintain a square visible at the center of the cross. Each pattern generated 30 repeated images across 8 mm, separated by 0.006 mm along the slow axis. The axial scan rate was 30 kHz. Subjects were aligned under 30 microwatts red light and imaged with 150 microwatts white light, which are conservatively safe and below maximum permissible exposures according to the American National Standard for Safe Use of Lasers (80). Alignment and imaging were performed with the same cross scan pattern.

### Human Subjects

Subjects were recruited from patient populations in general ophthalmology and optometry at the NYU Langone Eye Center. Sixty healthy human subjects without evidence of retinal disease (24-75 years old, 36 females, 8 blacks, 17 Asians, 30 whites, 4 unknown, and 1 multiracial, 102 eyes) were imaged. One 47-year-old female subject had neuromyelitis optica but denied any history of optic neuritis within the past 10 years. Seven glaucoma suspects were included. All procedures were reviewed and approved by the institutional review board at NYU Langone Hospital. Patients deemed eligible for participation were informed about the study via phone, secure email, or Epic message. An Eye Center delegated study team member then followed up with patients to explain the study. If a patient was interested, an appointment with the research team to conduct the consent process and study imaging was arranged. To ensure confidentiality, interested subjects were consented in secluded areas of the clinic. During the consent process, they were asked to provide basic information such as their age and history of eye disease. Extensive discussion of risks and possible benefits of participation was provided to the participants and their families. The study team members explained the research study to the participant and answered questions. The participant was asked to read and review the document before signing. A copy of the signed informed consent document was given to the participants for their records. The consent process, including the name of the individual obtaining consent, was thoroughly documented in the subject’s research record, and any alteration to the standard consent process was documented.

### Inclusion/Exclusion Criteria

Inclusion criteria for the study were: 1) between 18 and 99 years of age, 2) no significant anterior segment disease or media opacities such as corneal conditions or lens opacities (i.e. cataracts) limiting reliable visualization of retina, 3) no prior history or evidence of intraocular surgery (other than cataract extraction), 4) no history or evidence of retinal pathology (no drusen or RPE changes related to AMD), 5) best-corrected visual acuity of 20/40 or better, 6) intraocular pressure of 25 mmHg or lower, 7) no cataract surgery within 3 months or capsulotomy within 6 weeks prior to the visit, 8) no history of conditions in either eye which might preclude pupil dilation.

Exclusion criteria for the study were: 1) fundus is not visible, 2) media are opaque, 3) unwilling or unable to participate in the study. Note that the strict inclusion criteria resulted in excluding some subjects in their 80s and 90s with drusen or pigmentary changes that were not part of an AMD diagnosis.

### Image Reconstruction and Flattening

A series of complex images (transverse position versus depth) were reconstructed assuming a uniform tissue refractive index of 1.4. Threefold complex Fourier-transform based axial upsampling, approximate flattening, transverse and axial motion-correction, and depth-dependent dispersion compensation (81) were applied independently to each set of horizontal and vertical scans. When applicable, all OCT images were flattened to BM at the sub-pixel level through 4 steps: 1. Coarse flattening based on manually labeling the hyperreflective peak at BM (**Figure 5**A-B). 2. Transverse averaging across every 20 adjacent axial scans (**Figure 5**C), followed by cubic fitting of RPE multiple scattering tails, excluding BM, to achieve a scatter-corrected BM profile. It is possible that this method also removed ‘true’ sources of reflectivity, including intercapillary pillars, in the choriocapillaris. Gaussian fitting of said BM profile provided BM peak depth (**Figure 5**D). The BM peak depth (**Figure 5**D) versus transverse position was next fitted with a 5^th^ order polynomial curve (**Figure 5**E). Images were re-flattened according to this fitted curve, this time on a sub-pixel level, via complex Fourier transform-based upsampling. 3. Step 2 was repeated with transverse averaging across 10 adjacent axial scans. 4. Finally, an additional axial motion correction of repeated frames along the slow axis based only on the outer retina was performed to ensure no residual motion (**Figure 5**G). After application of the above procedures, outer retinal images show a flat BM (**Figure 2**C, **Figure 5**F), which can be partitioned into sub-regions (**Figure 5**H). In separate validation experiments, flattening was also applied to the IS/OS junction based on inner IS/OS boundary (49), which served as a negative control for age-related changes (**Fig. S**5). In the foveola, the sharp variations in band 2 (IS / OS) near the central bouquet result in some blurring of outer retinal anatomy as it is not possible to align both BM and IS / OS in foveally centered and nearly-centered frames.

**Figure 5.**
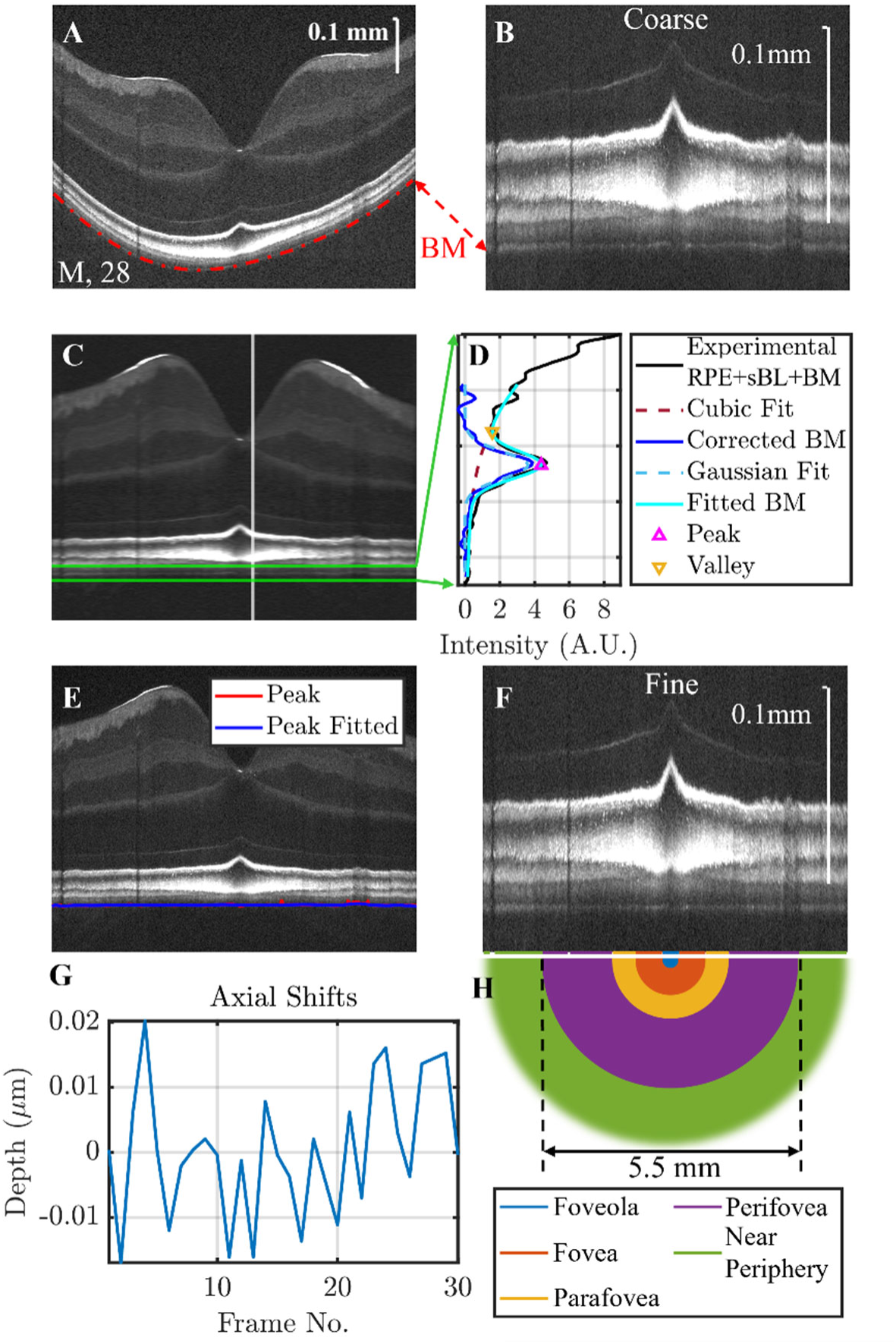
Image processing pipeline for a 28-year-old Asian male. All OCT images are in log scale. (A) OCT image with hyperreflective BM is labeled manually (red dash dot line). (B) Outer retina is coarsely flattened based on the label in (A). (C) OCT image is transversely averaged every 20 adjacent axial scans, with a designated depth range for BM peak localization range (green) and one example axial profile (white). (D) For each axial profile (black), a cubic fit depicts the RPE tail (brown dashed line) and a Gaussian fit depicts RPE scatter-corrected BM profile (blue dashed line). The BM peak depth is defined from the fitted BM profile (cyan). (E) BM peaks and fit versus transverse position. (F) Outer retina after the first round of subpixel level fine flattening, after which, a second round of subpixel level fine flattening was conducted with a transverse averaging size of 10 axial scans. (G) An additional motion correction along the slow axis based only on the outer retina shows very little residual motion. (H) Color-coded concentric half-circles indicate different eccentric regions: foveola (0.35 mm), fovea (1.5 mm, doesn’t include foveola unconventionally), parafovea (2.5 mm) and perifovea (5.5 mm), and the rest were classified as near periphery. Regions under blood vessels were excluded as needed from each sub-region.

Each OCT image was divided into 5 mutually exclusive concentric sub-regions (82): foveola, fovea, parafovea and perifovea with 0.35 mm, 1.5 mm, 2.5 mm, 5.5 mm diameter ranges around the foveal center respectively, and the rest were classified as near periphery (**Figure 4**C, **Figure 5**H). Note that the near periphery was not uniformly sampled in all eyes. For BM an average profile was created for each sub-region and each scan direction (H and V), then analyzed as described in the next section. For RPE+sBL, thickness was simply averaged over each sub-region and scan direction without radial weighting to account for polar coordinate geometry of the macular photoreceptor distribution.

### BM Morphometry

Based on physical understanding of light propagation through highly scattering melanosome and melanolipofuscin organelles in the RPE (47), a method was developed to quantify the thickness of BM (44). Essentially, the multiple scattering contributions from the RPE were removed by fitting with a third-order polynomial, leaving a corrected BM profile. This approach was validated with a Monte Carlo model of light propagation (47), and shown to reduce the coefficient of variation of BM measurements *in vivo* (44). This correction approach yielded a more mirror-symmetric, i.e. less skewed, BM profile which was more amenable to fitting with a Gaussian function. In instances where the BM peak was not visible, meaning no visible signature of BM in sub-region profiles meaning proper fitting ranges cannot be chosen (4.1% incidence), both BM thickness and contrast were marked as invalid, and not included in morphometric analysis. The incidence of indistinguishable BM was statistically analyzed.

The full-width-at-half maximum (FWHM) of a Gaussian fit to the corrected BM intensity profile (cyan dashed profiles in Fig. S1) was taken as BM thickness. To cross-check this method, the band 2 widths (FWHM) were quantified across different age groups. Band 2 inner half width is stable with age (**Fig. S5**).

The contrast was defined as the intensity ratio of the BM peak to the basal RPE+sBL valley (Figure. S4. A, B). Typically, in >90% of the profiles, the BM peak could be taken as the maximum of the profile (which also corresponded to a zero of the first derivative). Sometimes (e.g. Fig. S1C), there was no local maximum to designate as the BM peak for the contrast calculation, yet a BM profile was still clearly visible. In such instances (4.3% incidence), the BM hyperreflection is shown as an inflection point in the continuing downward of RPE tail (Fig. S1C), and contrast was less than 1. The depth of the local minimum derivative magnitude was employed as the “BM peak” depth. In these instances, after RPE tail correction, the depth at which the intensity reached 10% of the “peak” was taken as the “valley” depth (Fig. S1C). BM contrast was then calculated as above.

### RPE+sBL Morphometry

RPE+sBL height is taken as the distance between the segmented boundaries RPE-Inner and BM-Inner (see **Figure 2**C). The segmentation approach was adopted from prior studies (44). To investigate reflectivity across sub-regions and/or populations, longitudinal reflectivity profiles were normalized and stretched from 0% at the inner boundary of the segmented RPE cell body to 100% at the inner boundary of segmented BM. Stretched intensity profiles were compared among different age groups, between white and non-white subjects, and across retinal sub-regions.

### Statistical Analysis

The association between age and BM thickness, RPE+sBL thickness, and BM versus outer RPE+sBL contrast was evaluated using generalized estimating equation (GEE) models that accounted for intra subject correlation arising from bilateral measurements and measurements for both horizontal and vertical scan orientations. Models were adjusted for scan orientation (vertical or horizontal) to address potential anisotropy effects. No significant differences were observed between left and right eyes for each measurement, therefore laterality was not included as a covariate. Multiple comparisons were accounted for by calculating p-values using the false discovery rate (FDR) using the multipletests function from the statsmodels package in python. Pearson correlation coefficients were computed to examine the relationship between BM thickness and RPE+sBL thickness within each eccentricity.

Associations between the presence of photoreceptor anomalies and retinal thickness across eccentricities were assessed using GEE models with an exchangeable working correlation structure to account for repeated measures within each participant (across eyes and orientations). These models were adjusted for age, scan orientation, and retinal eccentricity, with an age-by-eccentricity interaction term to assess eccentricity specific age effects.

Regional associations between photoreceptor anomalies and retinal structural metrics were evaluated using GEE models, accounting for within-subject correlation due to inclusion of both eyes. Separate models were fit for retinal sub-regions with near periphery excluded (only one incident of a photoreceptor anomaly was observed in this sub-region) and outcome (BM thickness, RPE–BL thickness, and contrast), with adjustment for scan orientation (X/Y) in regions where orientation effects were present (perifovea and near periphery). FDR correction was applied within each outcome group to account for multiple regional comparisons.

## Acknowledgments

We acknowledge support from National Institutes of Health (EY031469, EY036192, EB035762, EB032840, OD038130); Research to Prevent Blindness (Stein Innovation Award and unrestricted departmental grant); and an NYU-KAIST Collaboration Award.

