## Supplementary Figures and Tables for "Visible Light Optical Coherence Tomography Reveals Aging at the Retinal Pigment Epithelium-Bruch’s Membrane Interface"

*Vivek Jay Srinivasan

**This PDF file includes:**

Figures S1 to S7

Table S1

SI References

### Figures

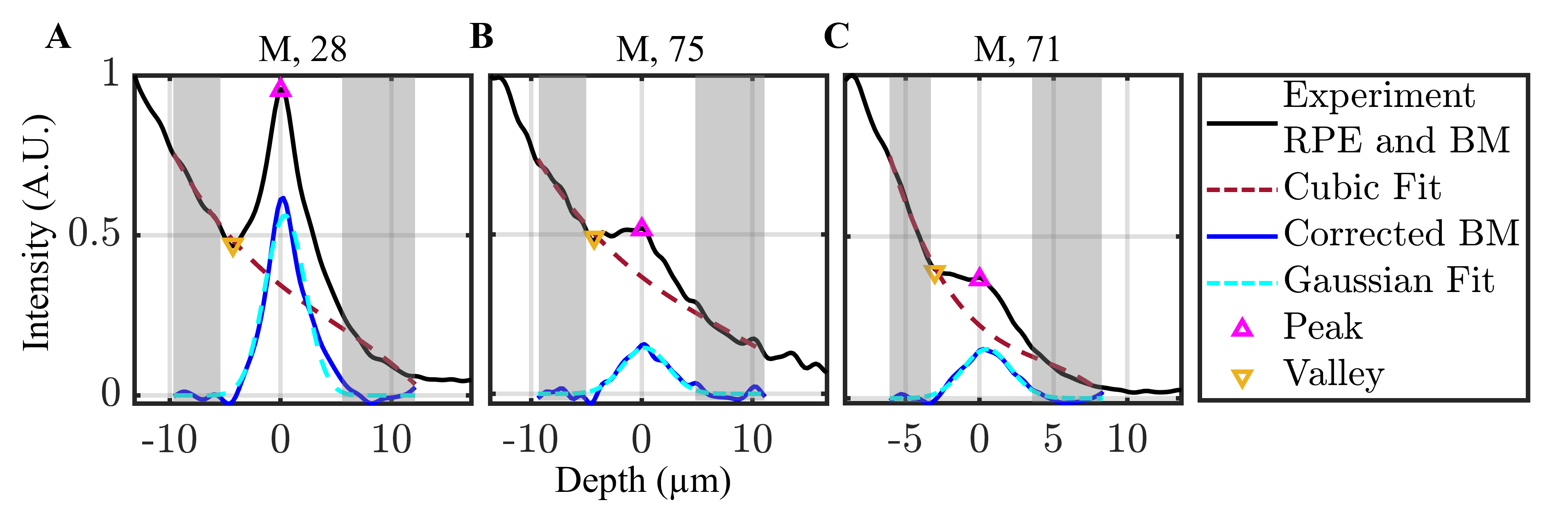

##### Fig. S1 BM thickness is determined as the width of a Gaussian fit (dashed cyan line) of the corrected BM profile (solid blue line) after cubic fitting (dashed brown line, ranges demarcated by shaded regions) of OCT intensity profile (solid black line) and subtraction to remove multiple scattering RPE tails. Contrast is defined as the ratio of BM ‘Peak’ (magenta upwards triangle) to RPE+sBL ‘Valley’ (orange downwards triangle). (A) 28-year-old male with a BM thickness of 2.56 micrometers and a BM contrast of 1.79; (B) 75-year-old male with a BM thickness of 2.88 micrometers and a BM contrast of 1.04; and (C) 71-year-old male with a BM thickness of 3.71 micrometers and a BM contrast of 0.95 are shown. Note that in the last profile, the “peak” was not detectable, so the local minimum derivative magnitude was employed as the “peak”. In these cases (4.3% incidence), and after RPE tail correction, the 10% point of the “peak” intensity was taken as the “valley”.

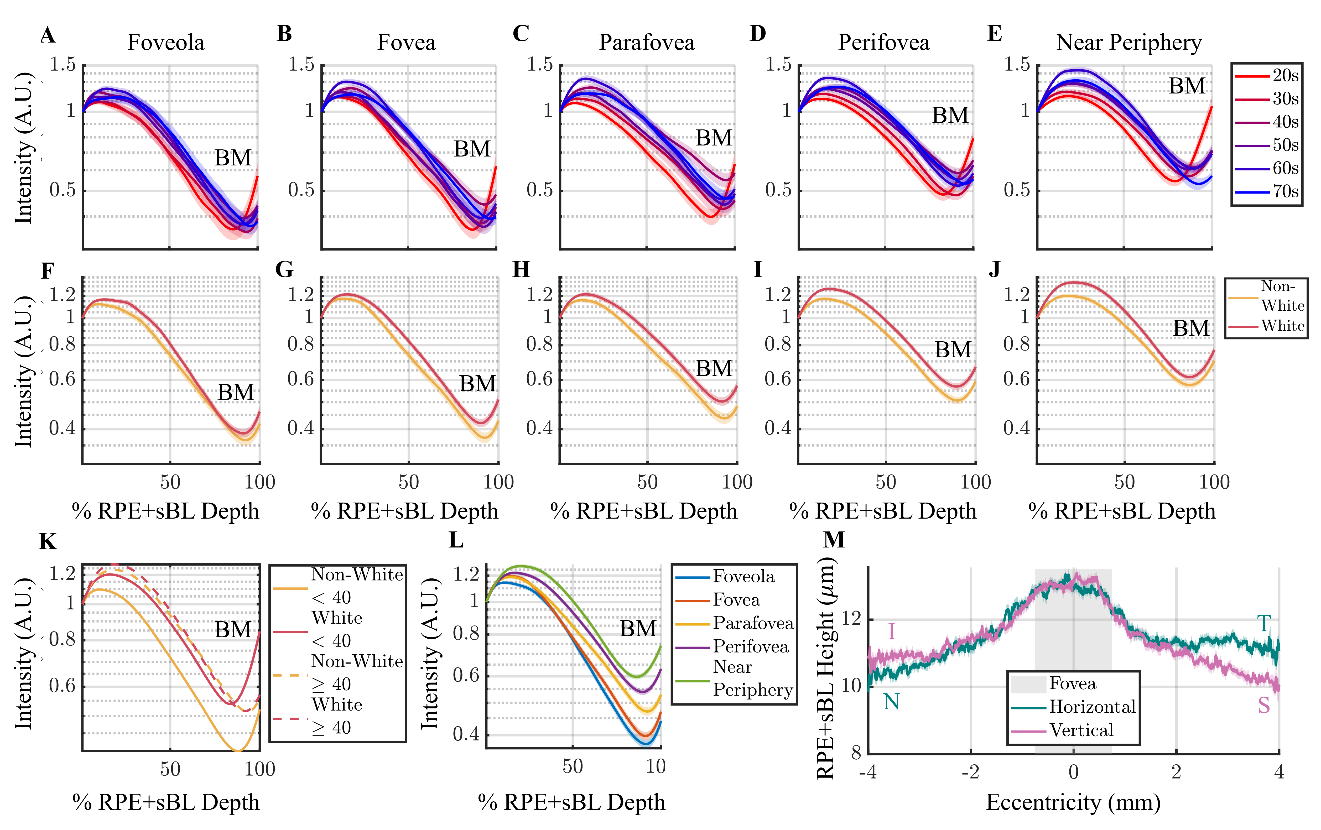

##### Fig. S2 Internal reflectivity of the retinal pigment epithelium (RPE) and the sub-RPE basal laminar space (together, RPE+sBL). An RPE+sBL depth of 0% is defined as the inner edge of the hyper-reflective RPE band. Intensity is normalized to 1 at this location. An RPE+sBL depth of 100% is defined as the inner edge of the hyper-reflective Bruch’s membrane (BM). (A-E) The most evident age-related changes are a diminishing of the BM peak with age and broadening of the apical reflectivity distribution into a plateau. The relative reflectivity in the nominal basal RPE+sBL region is diminished in older subjects, though the relationship with age is not monotonic and basal reflectivity is complicated by the more prominent BM peak in younger subjects. (F-J) The comparison between whites (49.9±15.2 years old) and non-whites (47.1±15.0 years old) shows a broadening of the apical-mid RPE reflectivity distribution for whites, particularly in the more eccentric sub-regions, similar to changes observed with age. (K) Reflectivity within RPE+sBL stratified by age and whites versus no-whites, averaged across sub-regions. Younger whites and non-whites (<40 years old, solid lines) show differing internal reflectivity; these differences disappear with age (≥40 year old, dashed lines). (L) Internal reflectivity across eccentricity mirrors replicates our previous results (44) from a younger cohort. (M) RPE+sBL height along horizontal (N: nasal, T: temporal) and vertical (I: inferior, S: superior) meridians show a decline towards superior and nasal sides, the latter of which was also shown histologically.

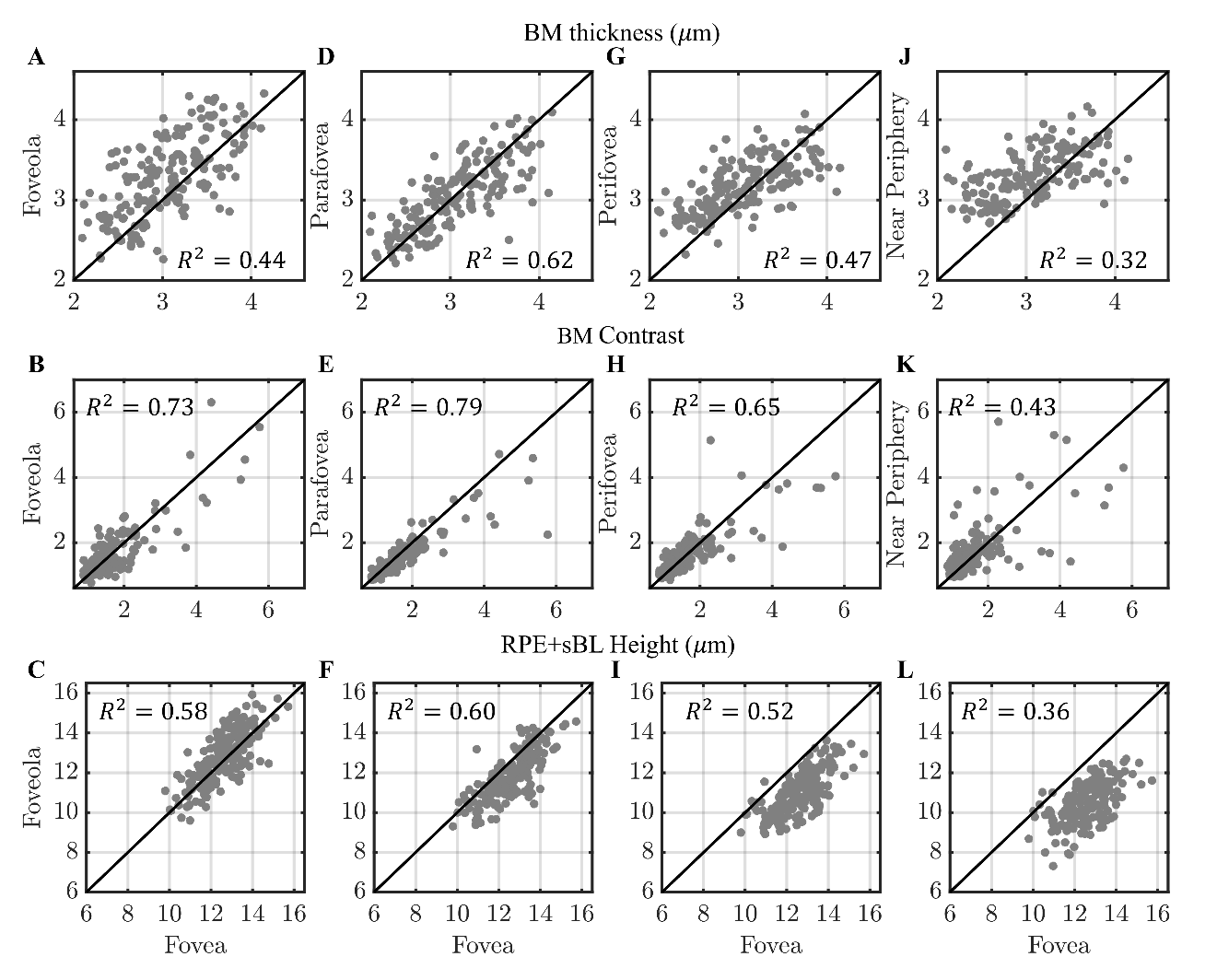

##### Fig. S3 Regional covariation of morphology. The foveal BM thickness (A, D, G, J), contrast (B, E, H, K), and RPE+sBL height (C, F, I, L) correlates with those in other sub-regions. A solid line of equality is shown for reference. Linear fits between each sub-regions show highest R^2^ between foveal and parafoveal sub-regions, and lowest R^2^ with near periphery. We find reduced performance of motion correction in the foveola, which may partially contribute to the high BM thickness deviation observed in that sub-region. In summary, morphology covaries across sub-regions, albeit with substantial variability, justifying the sub-regional analysis presented in this work.

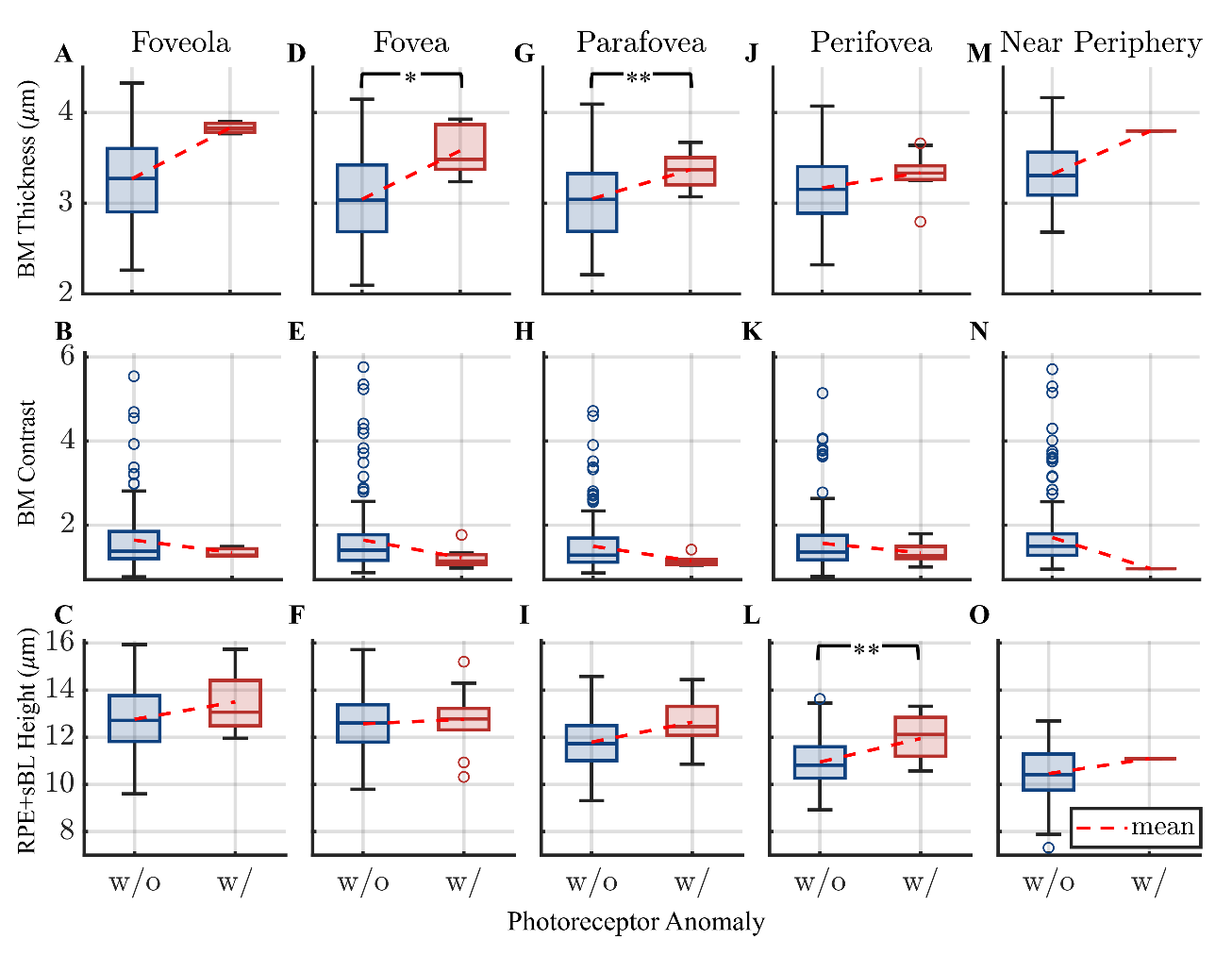

##### Fig. S4 Box-and-whisker plots for BM thickness, BM contrast, and RPE+sBL height, contingent on the presence or absence of a photoreceptor anomaly for foveola (A-C), fovea (D-F), parafovea (G-I), perifovea (J-L), and near periphery (M-O). Where a photoreceptor anomaly is present, BM is thicker, and RPE+sBL trends taller (BM thickness of foveal and parafoveal sub-regions, and RPE+sBL height in perifovea are significant in the GEE model. *p<0.05, **p<0.005). Though BM contrast trends lower with a photoreceptor anomaly present, comparisons are not statistically significant. Note that only one photoreceptor anomaly was found in the near periphery. Full p-values are shown in Table S1.

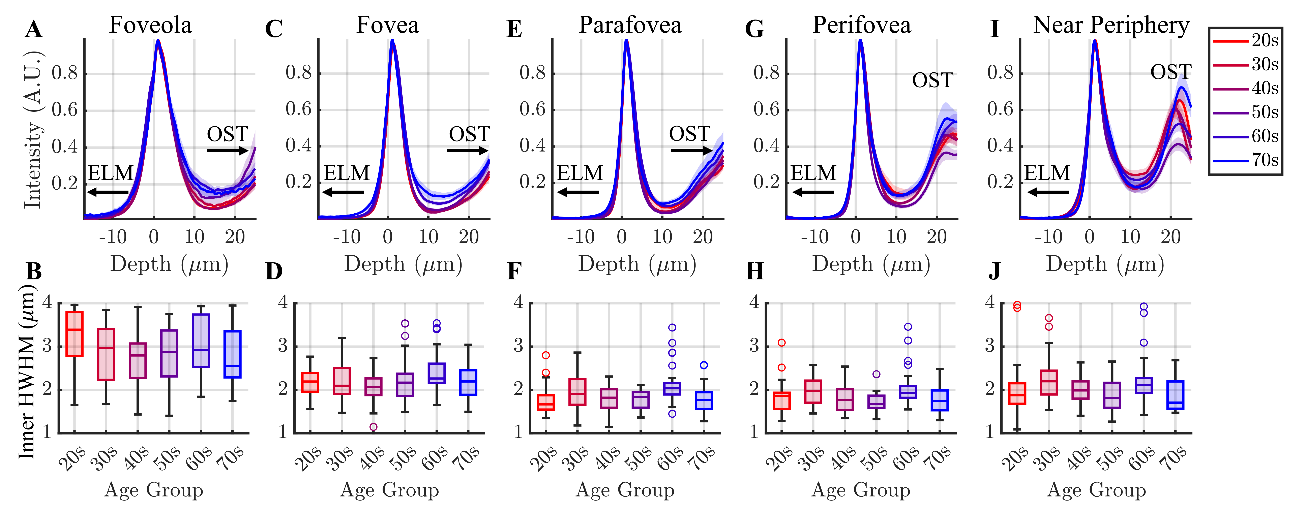

##### Fig. S5 Photoreceptor band 2, inner segment / outer segment junction (IS/OS) or ellipsoid zone (EZ), negative control. Band 2 profiles (A,C,E,G,I) and inner half widths (B,D,F,H,J, defined from the inner half-maximum to the peak) of six different age groups show that band 2 width is stable with age.

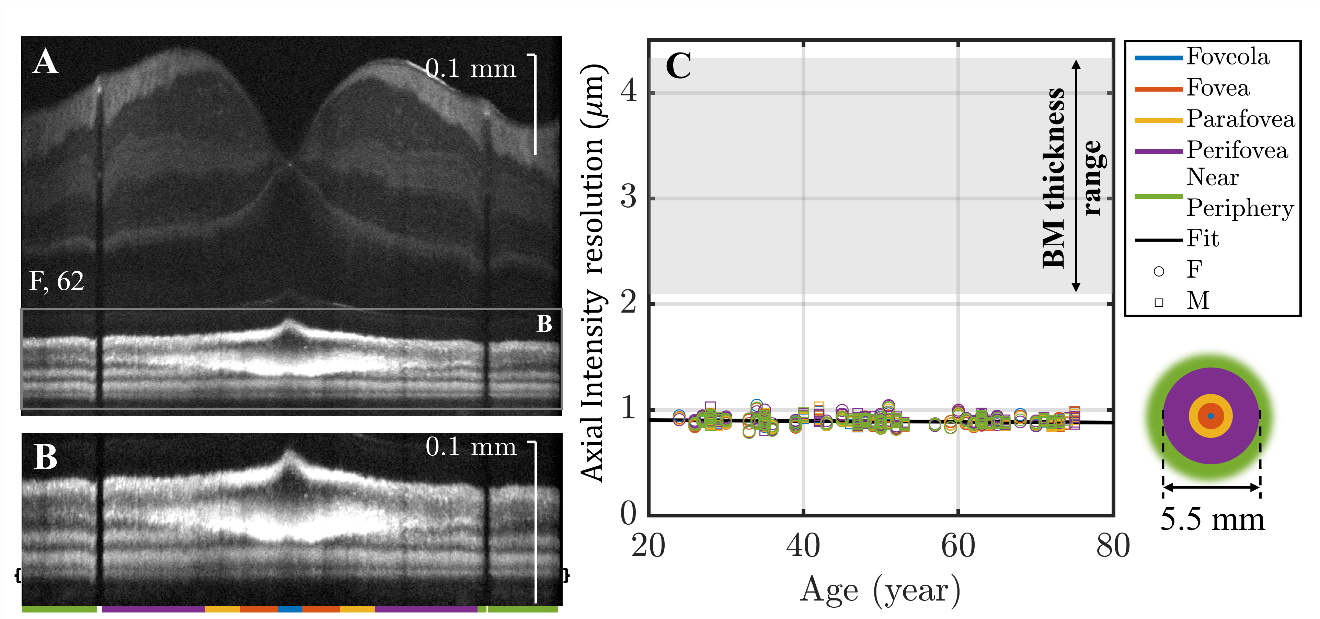

##### Fig. S6. Transform-limited visible light OCT depth resolution at the level of basal RPE+sBL and BM does not depend on age. Note that the incoherent intensity point spread function is analyzed due to averaging of images with independent speckles. (A) Retina and (B) outer retinal zoom of a 62-year-old white female subject, showing 5 sub-regions and the basal RPE+sBL+BM area (brackets) for achievable transform-limited axial resolution calculation. OCT images are in log scale. (C) Achievable transform-limited incoherent OCT resolution (circles and squares), based on the spectrum from basal RPE+sBL and BM, is significantly finer than range of measured BM thicknesses (brackets). Interestingly, the axial resolution in micrometers becomes finer, not coarser, with age (in years) as *∆z= 0.91 – (0.00034 µm/yr)×Age*. These results show that changes in spectral width cannot explain measured thickening of BM with age.

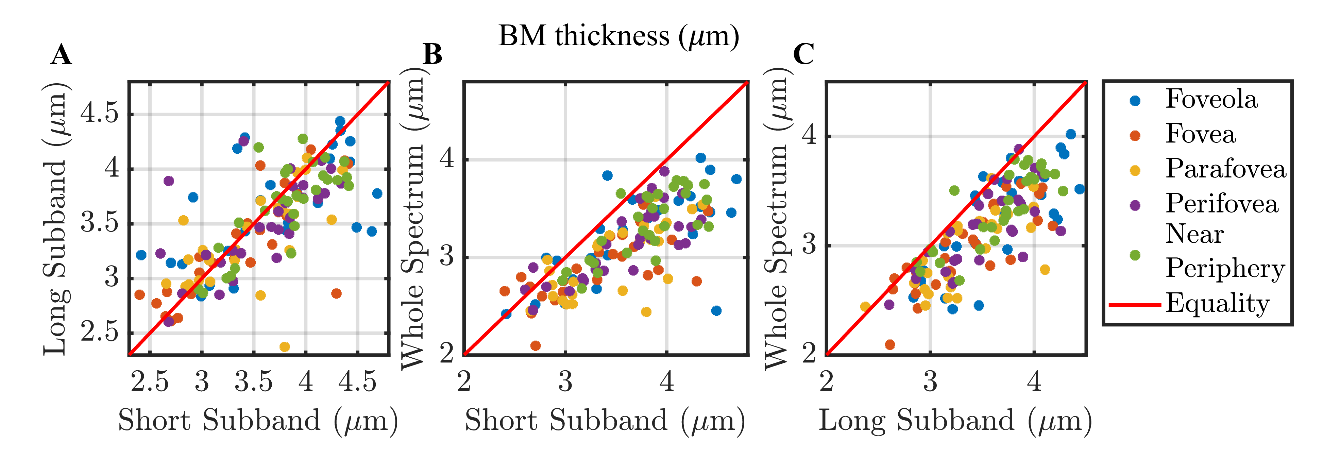

##### Fig. S7. In 9 representative subjects (6 female, 49.11±16.63 years old) BM thickness from long (centered at 527 nm) and short (centered at 579 nm) subbands with the same theoretical depth resolution (0.96 micrometers intensity FWHM) are identical (A). This finding argues against wavelength-dependent effects like scattering creating a systematic bias for BM thickness for this cohort. (B-C) Subband BM thicknesses are generally thicker than the BM thickness from the whole spectrum (0.71 micrometers intensity FWHM) indicating that resolution matters for BM thickness quantification.

| Model | GEE | | | |
| --- | --- | --- | --- | --- |
| Measurement | Foveola | Fovea | Parafovea | Perifovea |
| BM Thickness (µm) | 0.093 | 0.034 | 0.0036 | 0.23 |
| BM Contrast | 0.57 | 0.57 | 0.41 | 0.37 |
| RPE+sBL Height (µm) | 0.71 | 0.88 | 0.71 | 0.0035 |

Table S1. False data rate (FDR) corrected p-values based on GEE models for prediction of photoreceptor anomalies per sub-region. Note that p-values were not calculated in the near periphery sub-region since only one incident of a photoreceptor anomaly was observed.
